# A Study on multiple isotherms of Cr^6+^ ions using *Brevibacillus brevis* US575 isolated from tannery effluents

**DOI:** 10.1101/2023.05.28.542649

**Authors:** Arghyadeep Bhattacharjee, Om Saswat Sahoo, Srabani Karmakar, Arup Kumar Mitra

**Affiliations:** Department of Biotechnology, NIT Durgapur, Durgapur- 713209, West Bengal, India; Department of Microbiology, Kingston College of Science, Barasat, Kolkata- 120, West Bengal, India; Department of Microbiology, St. Xavier’s College Kolkata, Kolkata-16, West Bengal, India

**Keywords:** *Brevibacillus*, Chromium, Adsorption, Biosorbents, Tannery

## Abstract

Various leather factories in West Bengal have resulted in an enormous amount of leather that is processed annually. Regular discharges of tannery effluents into land and open water have led to significant soil and water pollution, with one of the most dangerous inorganic pollutants being chromium (Cr). This study aims to recover the most harmful components from industrial water using efficient biosorbents. *Brevibacillus brevis* US575 has been initially found in tannery effluents, where it has a high tolerance level to Cr^6+^ ions. The Cr(VI) adsorbed from the solution in aqueous phase during the course of the 60-minute contact period in this experiment was nearly 74%. Studies on the concentration of biomass, pH of the medium, and the starting concentration of metal ions have also been seen to affect the rate of biosorption. According to the desorption investigation, 1 M HCl outperformed all other concentrations of HCl, NaOH and pure water. Highest capacity of adsorption of the bio-adsorbent was calculated using the Langmuir model. The monolayer adsorption process was determined, and since the Freundlich model’s 1/n value fell inside 1, favourable adsorption has been postulated. According to the results of this study, the bacterium isolated from tannery wastewater was found to be the best alternative and could be used to create plans for using biosorption to combat current environmental pollution.

## Introduction

For the past few decades, urbanization and anthropogenic activities have increased massively, that leads to generation of life-threatening pollutants polluting the biodiversity, bioaccumulation and stability of the environment [1]. Inorganic metals like magnesium, manganese, calcium, nickel, chromium, sodium, and zinc are relatively less toxic and are needed for the redox and metabolic reactions of biosystems. However, heavy metals like chromium, mercury, lead and aluminium do not possess any biological role, and retain their toxicity to the society and destroys the ecological balance [2]. A similar response is observed when untreated tannery wastewater comprising toxic contaminants and heavy metals, especially chromium (Cr) is discharged [3,4]. Though, Cr(III) is useful to humans as it is associated with glucose and fat metabolism, Cr(VI) has been reported to be a health hazard, causing respiratory tract problems, and sometimes leading to lung tumors and gastric cancers [5]. Consistent exposure of Cr(VI) has also been shown to exert damage to DNA, and shorten life survivals of lower aquatic animals and plant life [6,7]. The release of toxic effluents from tannery industries, metal processing industries, preservative industries elevates chromium associated environmental hazards. Although different physical and chemical techniques like chemical precipitation, solvent partitioning and ion exchange chromatography have been utilized for the removal of toxic environmental contaminants, however these methods are not cost effective and require high energy. Though tannery water treatment, involving physiochemical methods like ion exchange, reverse osmosis and diffusion can be practiced, however biological wastewater treatment is favourable [8]. Bioremediation mediated by microorganisms and different microbial enzymes is also practiced to transform this toxic waste material into its comparatively lesser injurious state [9,10,11]. Being eco-friendly and cost competitive, it aids in revivality of the environment and maintains the ecological balance as well [12,13]. In our previous study, the role and efficacy of *Brevibacillus brevis* US575 in combatting Cr pollution has been well investigated. Also, the bacteria have been found to degrade not only chromium but also produces different industrially important enzymes like protease, lipase and keratinase. But as the demand is huge, different methods are also adopted to curtail these environment related issues [14]. In specific, biosorption possesses itself to be a common yet prominent and economical method to remove metal ions present in industrial effluents before release into waterbodies, as it includes biological sources like algae, bacteria, fungi etc [15,16,17]. Taking these facts into consideration, our study focuses on determining the efficiency of the biomass generated from bacteria isolated from tannery effluents as well as its underlying mechanism for the purpose of adsorbing toxic chromium ions.

## Materials and Methods

### Previous works

Study by Bhattacharjee et al. showed the proficiency of *Brevibacillus brevis* US575 in reducing Cr^6+^ to Cr^3+^ [14]. The bacteria isolated from water samples were subjected to serial dilution technique in dilutions ranging from 10^−1^ to 10^−4^. 0.1 ml of each sample was cultured in freshly prepared nutrient agar medium incorporated with 100-500 mg/L chromium, and were incubated overnight at 37ᵒC. All the four colonies were taken and sub-cultured separately to obtain further pure cultures. Results by Bhattacharjee et al., have also demonstrated the ability of *Brevibacillus brevis* US575 in degrading leather [14]. Thus this isolate was taken for preparing biomass and to understand its adsorption isotherms, amongst which the authors have adopted the Langmuir and Freundlich isotherms.

### Batch Adsorption Experiments

A mixture was prepared by taking 0.1 g biomass with 100 ml of metal solution at different concentrations (100, 500, 1000 mg/L) and was kept in a continuous stirring condition overnight. A definite amount of metal solution was collected at certain point of time and the solution pH was kept neutrality. At the end, every sample was filtered using a filter paper followed by measuring the remaining concentration of metal ions using atomic absorption spectrophotometer (AAS – Perkin Elmer (Analyst 200)). The experiments were performed in triplicates and the data was presented in mean and ±standard deviation (SD) [18]. The removal efficiency percentage was calculated as per standardised formula as described elsewhere.

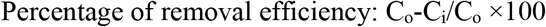

### Adsorption Isotherm study

Adsorption isotherm indicates the relationship between the adsorbate and the adsorbant surface, to which the adsorbate is adsorbed at equilibrium. Several models are well-standardized to study the adsorption isotherms of the adsorbate. In our study, two widely used models have been adopted, namely Langmuir and Freundlich isotherms.

### Desorption studies

An experiment was performed to recover the metal ions using 0.1 g of the biomass of the bacterium and 100 ml of 0.1 M of HCl and NaOH and was kept for a contact time period of 6–8 hr. Later, the same was recovered through the process of centrifugation at 8000 rpm for 12 min. The supernatant was filtered and stored at 4◦C until further experimental analysis. AAS analysis of the stored samples was performed to estimate the metal ion concentration [19]. The removal efficiency percentage was calculated as per standardised formula as described elsewhere.

## Results and Discussion

### Metal biosorption is dependent on contact time

The effect of contact time at different time intervals of 10, 20, 30, 40, 50, 60, 70, 80, 90, 100 mins was investgated. Our study indicates that highest metal absorption of nearly 78% was found at a time interval of 60 mins [Fig 1]. Also, no remarkable changes in the biosorption efficiency has been observed with increase in the time of contact. Ramachandran et.al., and his group reported that maximum uptake of metal was observed at 60 mins using soil bacteria *Bacillus amyloliquefaciens* that highly correlates with our findings [20]. However in another study using microalga *Chlamydomonas* sp., nearly 90% biosorption has been observed in 30 mins of contact time. [21]. Although various studies have have highlighted much more longer contact time to get maximum biosorption, studies by Zhang et.al showed that using a contact time of 60 mins using Bacillus subtilis, nearly 90% biosorption has been observed [22]. It is to be noted that longer contact time does not display significant changes changes in biosorption efficiency. Generally during longer time period, the surface area gets depleted hindering the binding the binding of metal ions on the surface.

**Figure 1.**
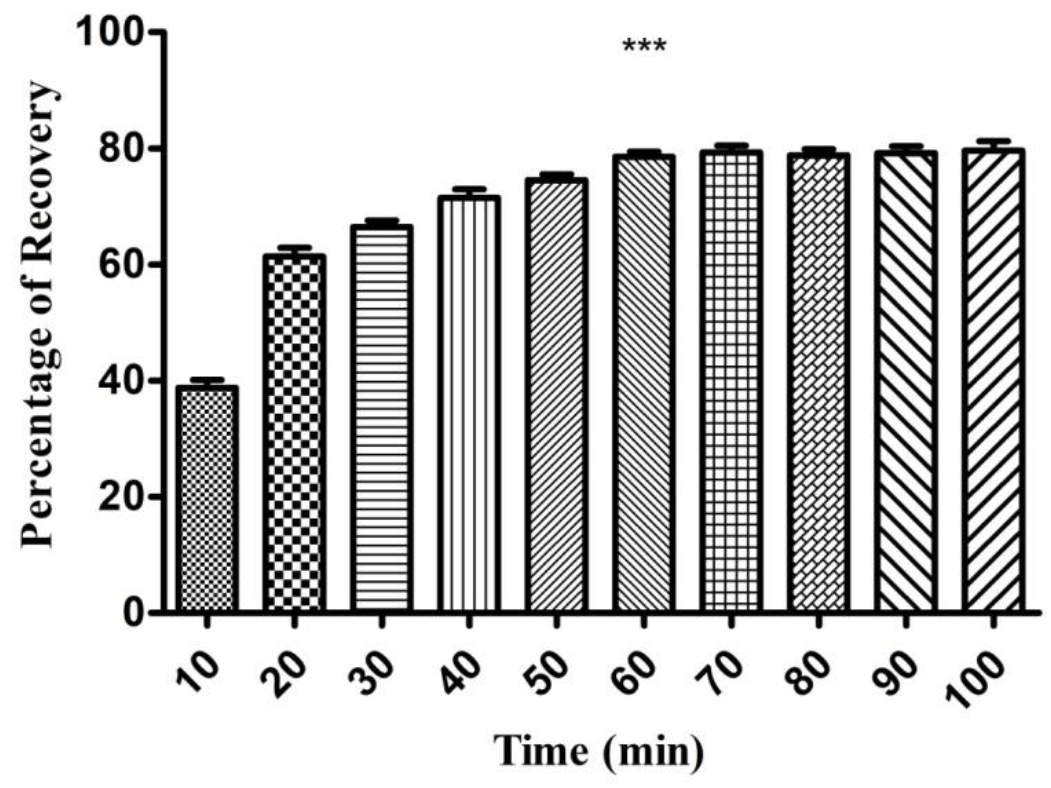
Graphical representation showing the change in metal biosorption in different time intervals. Data represented in the bar diagram showing mean ± SD. ***p < 0.0001, as determined by one-way ANOVA. SD: Standard deviation.

### The metal biosorption efficiency is dependent on the concentration of biomass

The effect of biomass at different concentrations ranging from (0.1-0.3) g on metal adsorption was investigated. The adsorbent dose was varied keeping the other parameters constant and the efficacy of *Brevibacillus brevis* US575 biomass in the absorption of Cr ions was studied. Our results highlighted that increasing the adsorbent quantity from 0.1 to 0.3 mg resulted in an increased metal intake capacity (Fig.2). On the contrary, no significant change in the metal intake capacity has been observed in further increase of biomass concentration (data not shown). It has been reported that maximum metal intake (absorption) was observed at a concentration of 0.3 g (Fig.2). Similar findings has been observed using *Bacillus amyloliquefaciens* where the metal adsorption capacity is maximum at 0.3 g [20]. Increase in the number of binding sites by the biomass strongly elevates the biosorption level to a certain extent. Various studies involving microorganisms such as *Spirulina* sp., *Spherotilus natans* etc. have documented that concentration of the biosorbent is directly proportional to metal intake capacity [23,24].

**Figure 2.**
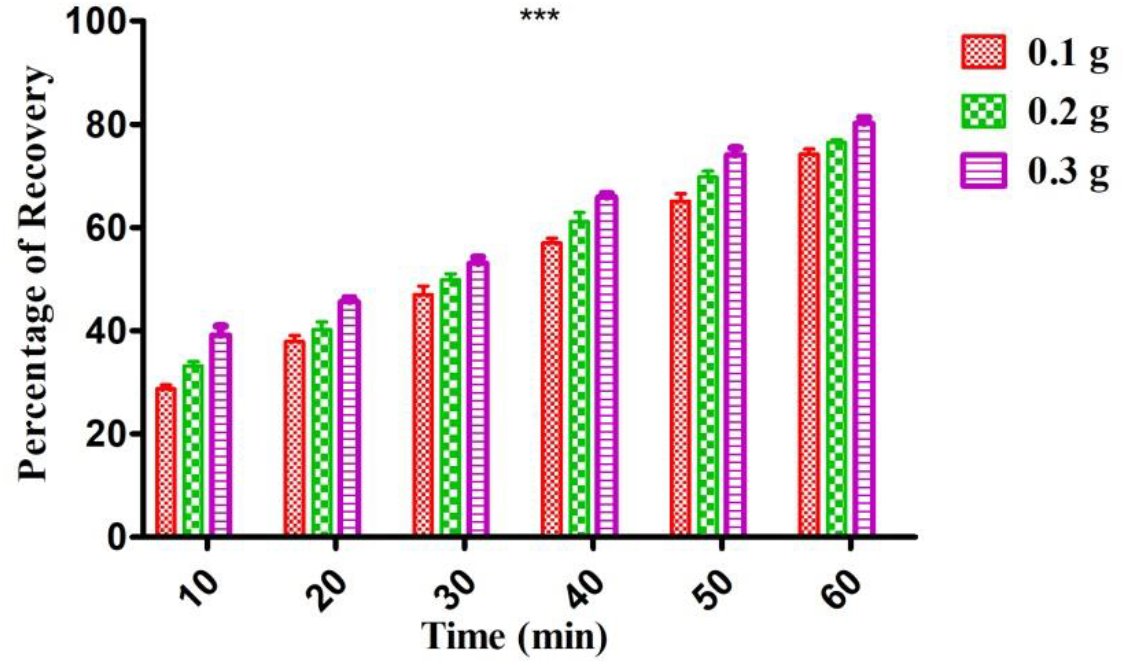
Graphical representation showing the effect of the concentration of biomass on metal biosorption at different time intervals. Data represented in the bar diagram showing mean ± SD. ***p < 0.0001, as determined by one-way ANOVA. SD: Standard deviation.

### The biosorption of heavy metal chromium is dependent on pH

The percentage of removal of heavy metal (chromium in our study) from the surrounding medium (aqueous solution) was studied at varying pH. In our study pH was varied from 3 to 9. Our study shows slight uptake of metal initially at pH 3 and this continued even after a contact time of 60 min. In pH 5 and 7, the metal uptake efficiency slightly increased at every time points. The efficiency of biosorption ranged between 15.8% to 61% at pH 3 to 7 at different time points (Fig. 3). On the contrary, the highest metal removal percentage of nearly 70% by the bacteria was found at pH 9 which showed that pH 9 is the optimum pH that will be considered for removal of heavy metal using the biomass of *Brevibacillus brevis* US575 (Fig. 3). The metal removal percentage at pH 9 ranged between 32.4% to 70.8% at different time intervals taken in our study. In a study by Tan et.al., it was shown that *Bacillu*s sp., CRB-B1 removes chromium at an optimum pH ranging between 6-8 [25]. In a similar study, *Oceanobacillus profundus* has been shown to exhibit maximum biosorption at a pH of 6 [26]. On the contrary, *Bacillus amyloliquefaciens* removes toxic chromium ions maximally at a neutral pH [20]. In low pH, the metal intake capcity is low owing to weak interaction between the metal and the surface of the biomass.

**Figure 3.**
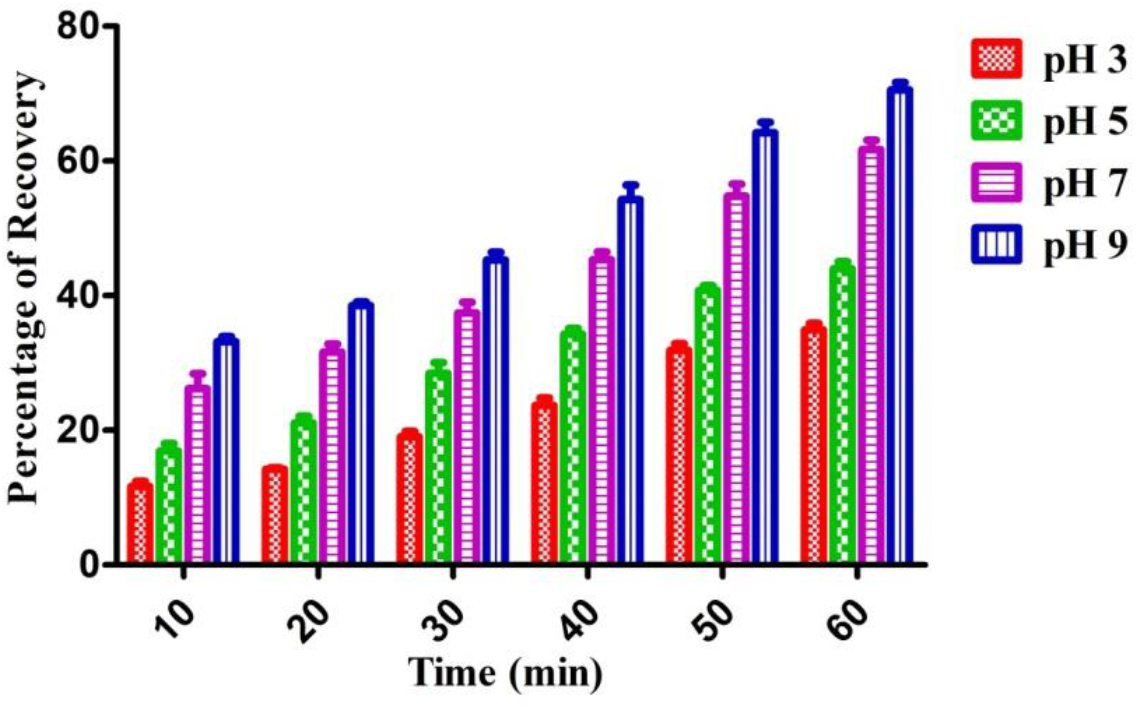
Bar graph showing the effect of the concentration of pH on heavy metal biosorption at different time points. Data represented in the bar diagram showing mean ± SD. ***p < 0.0001, as determined by one-way ANOVA. SD: Standard deviation.

### Study of Adsorption isotherms

The two models were plotted in their respective plots and were presented in Fig. 4A & 4B. It has been observed from our study that the experimental data obtained was best fitted for the metal ion used in our study. The capacity of the biosorbent to adsorb maximally (q_max_ (mg/g)) was 46.35 mg/g as obtained from the Langmuir model. Table 1 & 2 shows that, in comparison to Langmuir (R^2^=0.94), the model of Freundlich has yielded a better fit of R^2^= 0.98. Also, it has been found that the mechanism of adsorption was of monolayer type. From the Freundlich model, it has also been indicated that the 1/n value is within 1 indicating a favourable adsorption process [27]. Likewise, the separation factor (RL) is also less than 1 showing a suitable adsorption.

**Table 1:**
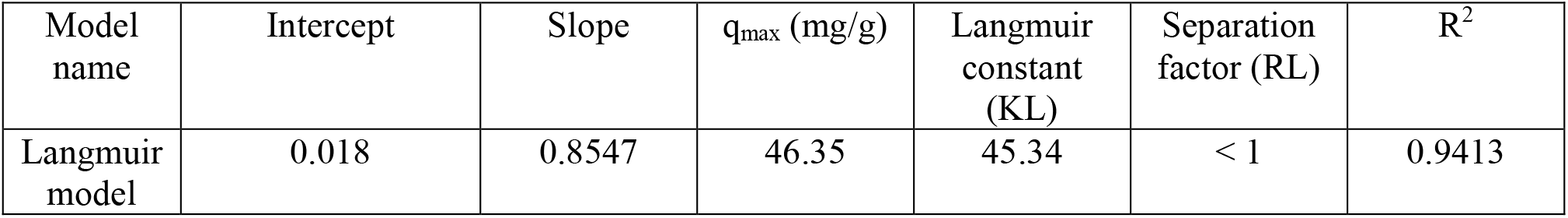
Langmuir model of biosorption isotherm for Cr^6+^ ions

| Model name | Intercept | Slope | q<sub>max</sub> (mg/g) | Langmuir constant (KL) | Separation factor (RL) | R<sup>2</sup> |
| --- | --- | --- | --- | --- | --- | --- |
| Langmuir model | 0.018 | 0.8547 | 46.35 | 45.34 | < 1 | 0.9413 |

**Table 2:** Freundlich model of biosorption isotherm for Cr^6+^ ions

| Model name | Intercept | Slope | 1/n | Freundlich adsorption constant (Kf) | R<sup>2</sup> |
| --- | --- | --- | --- | --- | --- |
| Freundlich model | 0.2769 | 0.7062 | 1.43 | 1.4 | 0.9871 |

**Figure 4.**
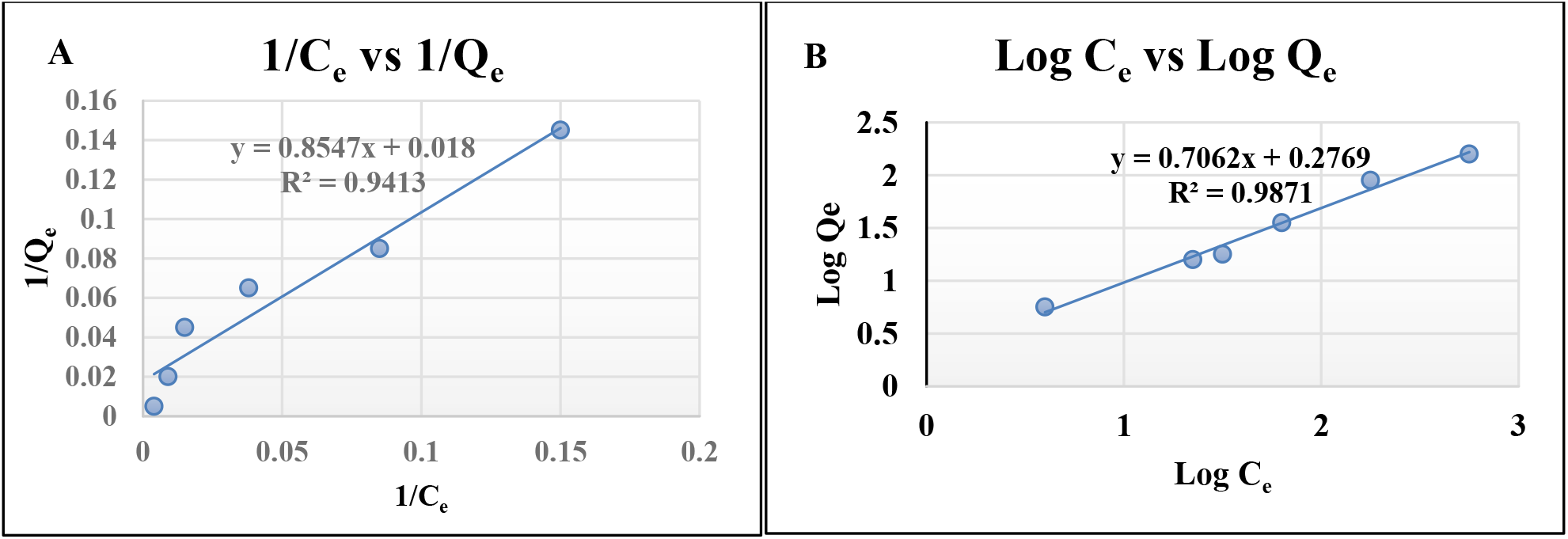
Graphical plots representing Adsorption isotherms of Cr^6+^ by (A) Langmuir model (B) Freundlich model

### Study on desorption by the bacterial biomass

The biosorbent derived from *Brevibacillus brevis* US575 that has been treated with 1 M hydrochloric acid showed nearly 84% recovery of heavy metal chromium in 60 minute (Fig.5). However, the same when regenerated with 1M NaOH and 0.5M HCl showed recovery percentage of 59% and 63% respectively at the maximum time point (60 minute) (Fig.5). Our study also highlighted that the same biosorbent is associated with low desorption efficacy when treated with deionized water and 0.5 M NaOH (Fig.5). A more or less similar result has been obtained by Govindam and his group when they used *Bacillus amyloliquefaciens* resulting in a metal recovery rate of nearly 90% [20]. A study reported from China using *Bacillus subtilis* showed nearly 94% desorption rate when the biomass waste have been treated with HCl for 1 hour [22].The proposed mechanism is that the positively charged cation in the metal ion interacts with the negatively charged functional group present on the surface of the biosorbent. Electrostatic interactions within the surface of the biosorbent were produced by the metal ion present in the waste water [28]. Studies involving heavy metals like Cd, Pb etc. have reported similar type of absorption mechanism [19]. Owing to this type of interaction mechanism, maximum desorption is achieved only using 1 M acid.

**Figure 5.**
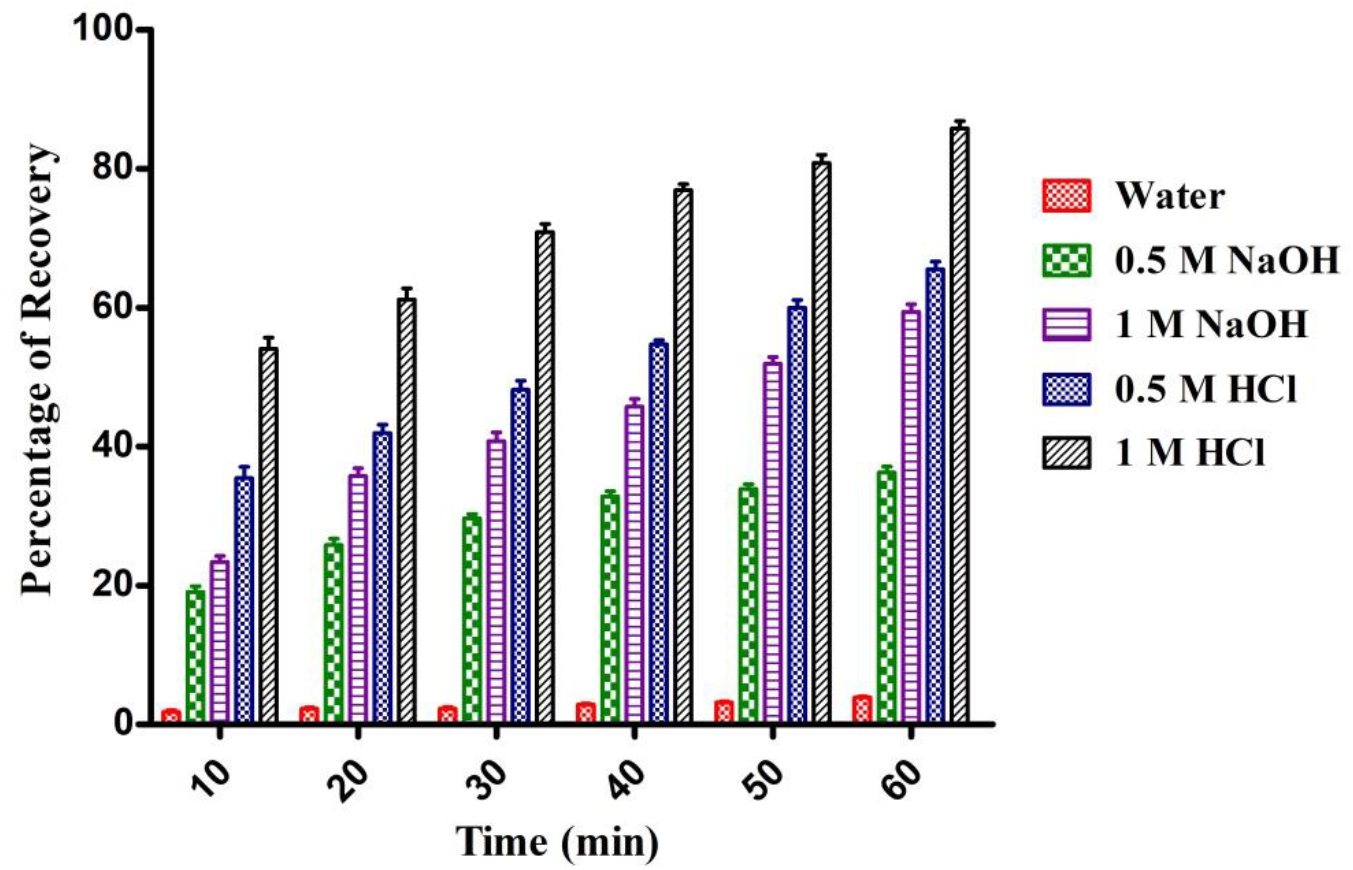
Graphical representation showing desorption studies of *Brevibacillus brevis* US575 biomass at different time points. Data represented in the bar diagram showing mean ± SD. ***p < 0.0001, as determined by one-way ANOVA. SD: Standard deviation.

## Conclusion

Remediation of industrial effluents, primarily toxic metals, is essential in terms of safeguarding both the environment and human health. In the present investigation, Cr^6+^ in the tannery effluents can be removed using biomass of *Brevibacillus brevis* US575. In addition, biosorption studies further revealed that the efficiency of biosorption was resisted by low pH and the time period of contact. Throughout our study, optimum biosorption was observed at a pH and contact period of 9 and 60 min respectively. However, to apply this process on a large scale, i.e. from lab to industries, further studies like chemical modifications of the biosorbents along with improvisation in biosorption parameters might be highly needed. These findings can be improvised by small-scale piloting and, can be translated on a large scale to clean up chromium concentrated water bodies, including tannery effluents before discharging the wastes into the environment.

## Acknowledgements

The authors express their deep gratitude of thanks to the Department of Microbiology, Kingston College of Science, Department of Microbiology, St. Xavier’s College, Kolkata, Department of Biotechnology, National Institute of Technology Durgapur. The authors are also deeply indebted to Rev. Dr. Dominic Savio, S.J., Principal and Rector, St. Xavier’s College (Autonomous), Kolkata, Dr. Srabani Karmakar, Principal, Kingston College of Science, Kolkata for their constant support and guidance throughout.

## Author contribution

The experimental design was performed collectively by all the authors. Preparation of reagents and samples along with data collection and analysis was done by Arghyadeep Bhattacharjee, Srabani Karmakar and Arup Kumar Mitra. The manuscript was written by Arghyadeep Bhattacharjee and Om Saswat Sahoo with inputs from all the authors Arghyadeep Bhattacharjee—Draft manuscript preparation, methodology, data collection and editing, Om Saswat Sahoo— manuscript editing, Srabani Karmakar—data interpretation and resources, Arup Kumar Mitra—conceptualization, investigation, data curation and overall supervision. All the authors read and approved the final manuscript.

## Declarations Ethics Approval

Not applicable

## Consent to Participate

Not applicable

## Consent to Publish

All the authors consent to the consideration for this manuscript for publication.

## Data Availability

Not applicable

## Availability of data and materials

Not applicable

